# Spatiotemporal Dynamics of Invariant Face Representations in the Human Brain

**DOI:** 10.1101/2025.06.03.657646

**Authors:** Amita Giri, Grace Smith, Cassia Manting, Katharina Dobs, Amir Adler, Dimitrios Pantazis

## Abstract

The human brain can effortlessly extract a familiar face’s age, gender, and identity despite dramatic changes in appearance, such as head orientation, lighting, or expression. Yet, the spatiotemporal dynamics underlying this ability, and how they depend on task demands, remain unclear. Here, we used multivariate decoding of magnetoencephalography (MEG) responses and source localization to characterize the emergence of invariant face representations. Human participants viewed natural images of highly familiar celebrities that systematically varied in viewpoint, gender, and age, while performing a one-back task on the identity or the image. Time-resolved decoding revealed that identity information emerged rapidly and became increasingly invariant to viewpoint over time. We observed a temporal hierarchy: view-specific identity information appeared at 64 ms, followed by mirror-invariant representations at 75 ms and fully view-invariant identity at 89 ms. Identity-invariant age and gender information emerged around the same time as view-invariant identity. Task demands modulated only late-stage identity and gender representations, suggesting that early face processing is predominantly feedforward. Source localization at peak decoding times showed consistent involvement of the occipital face area (OFA) and fusiform face area (FFA), with stronger identity and age signals than gender. Our findings reveal the spatiotemporal dynamics by which the brain extracts view-invariant identity from familiar faces, suggest that age and gender are processed in parallel, and show that task demands modulate later processing stages. Together, these results offer new constraints on computational models of face perception.

## 1 Introduction

Recognizing familiar faces is essential for human connection and communication. We can identify someone we know in a wide range of contexts—spotting a friend in a crowd, recognizing a celebrity on screen, or recalling a relative after years apart. This remarkable ability reflects a fundamental computational challenge: recognizing the same individual across highly variable visual input, such as unusual angles, variable lighting, or changes in expression or hairstyle. A key component of this challenge is achieving *view invariance*—the ability to recognize a person regardless of head orientation. Moreover, we do not just recognize who someone is; we also rapidly extract other socially relevant attributes such as age and gender. Yet, the spatiotemporal neural dynamics underlying the extraction of identity, age, and gender from familiar faces across such varied conditions remain incompletely understood.

Prior research has identified a distributed network of brain regions involved in face perception, including the fusiform face area (FFA), occipital face area (OFA), and posterior superior temporal sulcus (STS) [1, 2]. Neural evidence using fMRI suggests a hierarchical progression from view-specific coding in early visual regions and OFA, to increasingly invariant representations in FFA and downstream areas such as the anterior temporal lobe and inferior frontal cortex [3–5]. However, fMRI lacks the temporal resolution needed to pinpoint when invariant identity representations emerge. High-temporal resolution studies using EEG and MEG have suggested that view invariance unfolds over time: early neural activity reflects head orientation, followed by mirror-symmetric responses, and later emergence of invariant representations [6, 7]. However, these studies typically did not decode view-invariant identity directly, possibly because they used unfamiliar faces.

In contrast, familiar face recognition is thought to recruit additional brain systems, including regions involved in theory of mind and memory retrieval [8], and is more robust to noise, changes in appearance, and degraded image quality than unfamiliar face recognition [9]. According to the familiarity advantage hypothesis, repeated exposure enables the formation of more robust, invariant representations of familiar individuals, possibly supported by top-down mechanisms that facilitate early visual processing [10]. Indeed, previous work found image-invariant identity representations for familiar, but not for unfamiliar faces [11]. However, it remains unclear whether and how this familiarity-driven robustness manifests in the temporal dynamics of view-invariant identity representations.

Beyond identity, the brain also extracts other facial dimensions such as age and gender—attributes that are critical for social perception and decision-making. Although it is often assumed that such features are processed early and rapidly, the temporal order in which identity, age, and gender are extracted remains unclear. Some studies suggest that age and gender are extracted earlier than identity because they rely on coarse visual features [11], while others report early modulation of identity by gender [12]. Discrepancies across findings may reflect differences in stimulus types, task demands, or familiarity of the faces used.

Despite extensive research on face processing, several key questions remain unanswered. First, how does the brain achieve view-invariant recognition of familiar faces over time? Second, in what temporal sequence are different facial dimensions—identity, age, and gender—extracted? And third, to what extent do task demands shape these neural representations across processing stages [13] Yet most prior studies rely on unfamiliar faces and overly standardized images with limited ecological validity [3, 6, 7, 14, 15]—conditions that may underestimate the complexity and robustness of real-world face perception.

To address these questions, we recorded high-density MEG while participants viewed naturalistic images of twelve well-known celebrities, systematically varying in age, gender, and head orientation. To manipulate task demands, participants performed one of two one-back tasks: detecting image-level repetitions or identity-level repetitions. Using time-resolved decoding and source localization, we tested three core hypotheses: (1) identity representations become increasingly invariant to head orientation over time; (2) identity, age, and gender emerge in a consistent temporal sequence; and (3) task demands modulate neural representations across the visual processing cascade. By combining ecologically valid stimuli with temporally precise neural measurements, our study provides new insights into how the brain extracts person-specific information under real-world conditions.

## 2 Results

### 2.1 Behavioral performance during MEG task

We recorded MEG data from 19 participants while they viewed random sequences of familiar faces. We chose 12 celebrities highly familiar to U.S. adults, who varied orthogonally in gender and age, resulting in an equal distribution of six women and six men, as well as an even split between younger and older individuals. A total of 15 images per celebrity were selected, consisting of three distinct images for each of five specified head views: direct, half left profile, full left profile, half right profile, and full right profile (Fig. 1a). Images for each identity were chosen to incorporate natural variability in various dimensions, such as hair color, hair style, or background.

**Fig. 1:**
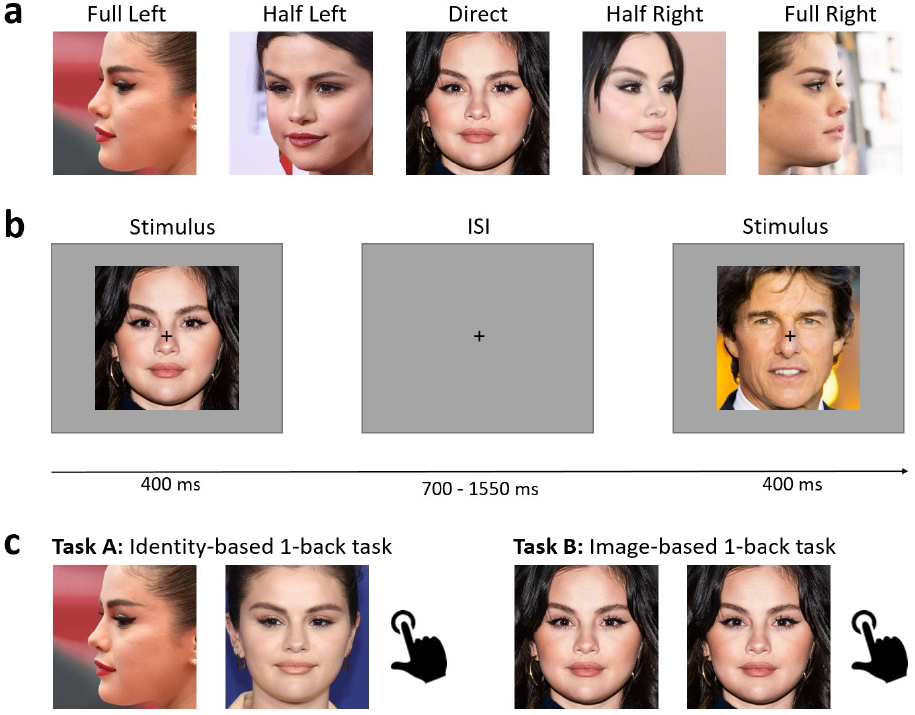
Experimental design. (a) A total of 15 images per celebrity were selected, consisting of three distinct images for each of the five head views: direct view, half left profile, full left profile, half right profile, and full right profile. (b) During the MEG experiment, subjects viewed a random sequence of familiar face images. Each trial started with the presentation of a face image for 0.4 s followed by a 0.7–1.55 s interstimulus interval. (c) To study task effects, participants conducted two tasks in different experimental sessions. In Task A, participants were instructed to press a button when the same person appeared consecutively, while in Task B, they responded when the same image appeared consecutively. The exact stimulus set is available at https://osf.io/eh54u/.

Participants viewed 10 repetitions of each of the 180 face images, each presented for 400 ms in individual trials (Fig. 1b). All images were presented in random order. To enhance face representations and study head view effects, participants were instructed to perform a one-back identity-based task (Task A), which required them to press a button with their right thumb when they saw the same person consecutively (Fig. 1c, left). Additionally, to investigate task influences on identity processing alongside two other facial attributes - age and gender - participants completed a separate session with a image-level task. In this one-back image-based task (Task B), participants were prompted to identify instances where the same image was presented consecutively (Fig. 1c, right). The order of the two experimental sessions was counter-balanced across participants.

Participants demonstrated heightened sensitivity to image repetition (mean sensitivity index d’ ± SEM: 4.72 ± 0.88) compared to identity repetition (mean sensitivity index d’ ± SEM: 3.73 ± 0.67). This result suggests that the identity-based task demanded a greater level of attention relative to the image-based task.

### 2.2 Time course of identity decoding invariant to head view

To unveil how humans perceive identity despite variations in head orientation, we conducted a time-resolved multivariate pattern analysis on the MEG signals collected during the identity-based one-back task. This analysis was conducted individually for each participant. At each time point, a support vector machine (SVM) classifier was employed to conduct binary classification between every pair of identities using the MEG sensor measurements. The classifier was trained using a 5-fold cross-validation procedure, where trial assignments to the folds were designed to yield outcomes sensitive to either non-invariant or invariant face representations concerning head view. Specifically, to capture the temporal dynamics of identity *non-invariant* to head view, MEG trials were randomly allocated to folds regardless of head view. Conversely, to isolate the temporal dynamics of identity *invariant* to head view, MEG trials sharing the same view among the five were grouped within the same fold. Thus, the classifier underwent training and testing on different views, enabling evaluation of its generalization across unseen head orientations. A pictorial representation of how different head views were assigned to folds is depicted in Fig. 2.

**Fig. 2:**
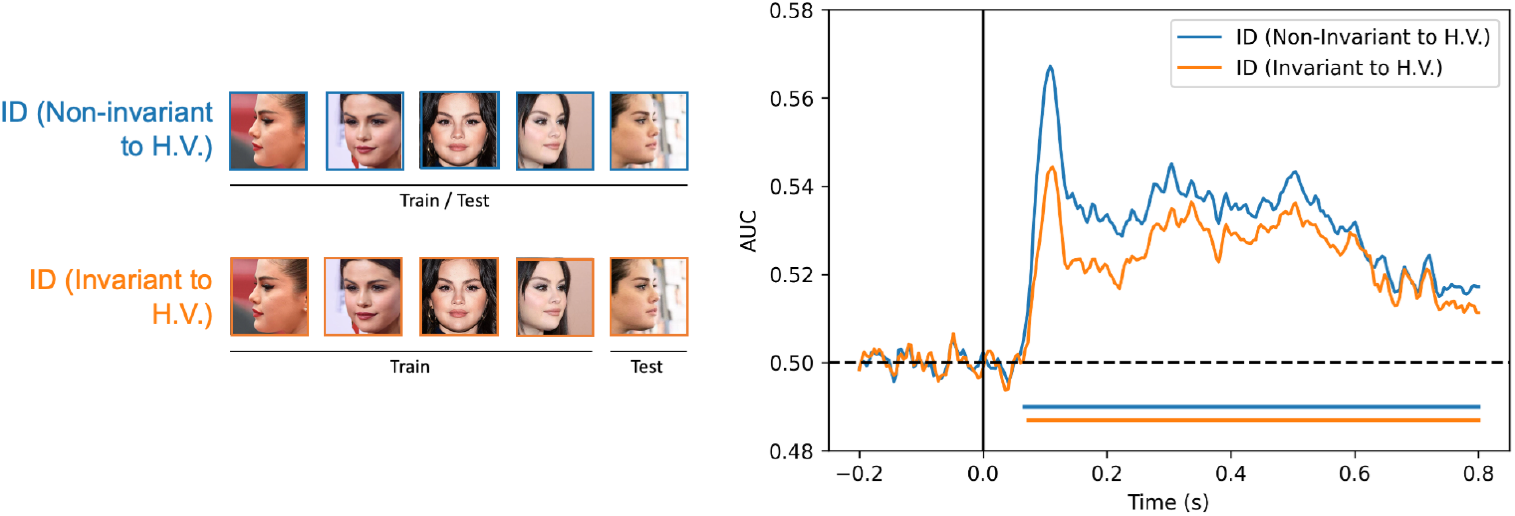
Time course of identity (ID) decoding non-invariant and invariant to head view. A classifier was trained to pairwise discriminate identities, utilizing either identical views for both training and testing sets (non-invariant), or varied views (invariant). Time 0 denotes stimulus onset. Lines below plots indicate significant time points using a cluster-based sign permutation test (cluster-defining threshold p *<* 0.05, and corrected significance level p *<* 0.05).

Classification results were averaged across participants and identity pairs, resulting in decoding time courses for both non-invariant and invariant conditions (Fig. 2). We found that identity representations non-invariant to head view generally exhibited greater robustness compared to their invariant counterparts, as indicated by a higher Area Under the Receiver Operating Characteristic Curve (AUC). Table 1 summarizes the onset and peak latencies of the decoding time courses, and Table 2 presents the corresponding statistical tests over latency differences (conditions 1 and 2). Notably, non-invariant representations emerged at 67 ms (95% confidence interval: 48 - 74 ms), while invariant representations appeared later at 73 ms (61 - 77 ms). The onset of non-invariant representations was significantly earlier than that of invariant representations (Δ = 6.01 ms, *P* = 0.028*), suggesting an early emergence of identity representations non-invariant to head view, followed by later transition to invariance with head orientation.

**Table 1:**
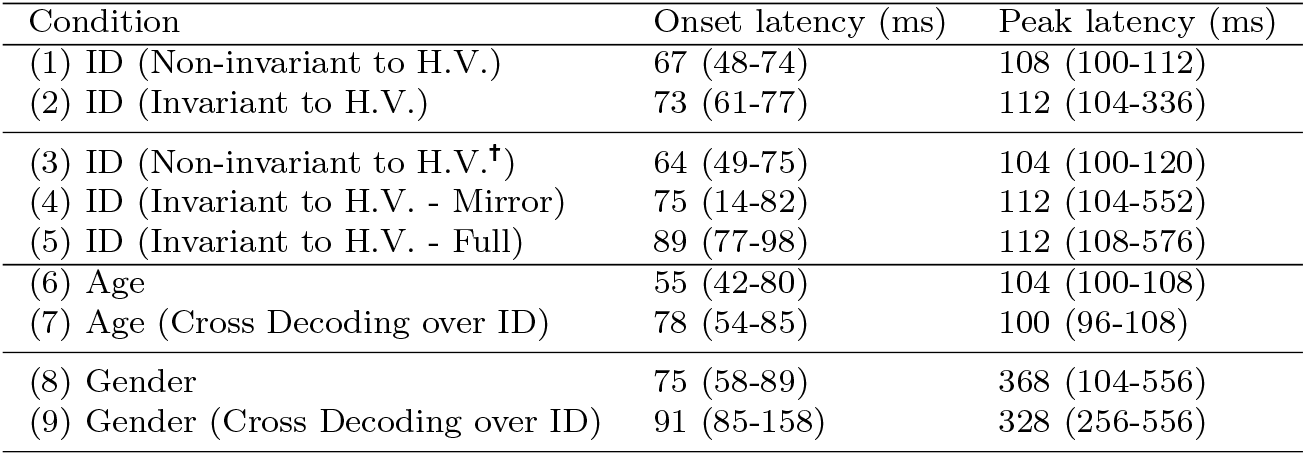
Peak and onset latency for different experimental conditions. Numbers in brackets indicate 95% confidence intervals. ^†^ Number of trials were matched

| Condition | Onset latency (ms) | Peak latency (ms) |
| --- | --- | --- |
| (1) ID (Non-invariant to H.V.) | 67 (48-74) | 108 (100-112) |
| (2) ID (Invariant to H.V.) | 73 (61-77) | 112 (104-336) |
| (3) ID (Non-invariant to H.V.<sup>†</sup>) | 64 (49-75) | 104 (100-120) |
| (4) ID (Invariant to H.V. - Mirror) | 75 (14-82) | 112 (104-552) |
| (5) ID (Invariant to H.V. - Full) | 89 (77-98) | 112 (108-576) |
| (6) Age | 55 (42-80) | 104 (100-108) |
| (7) Age (Cross Decoding over ID) | 78 (54-85) | 100 (96-108) |
| (8) Gender | 75 (58-89) | 368 (104-556) |
| (9) Gender (Cross Decoding over ID) | 91 (85-158) | 328 (256-556) |

**Table 2:** P-values for peak and onset latency differences between the different conditions listed in Table 1. Statistical tests were one-sided or two-sided t-tests where appropriate. P-values were corrected for multiple comparisons using false discovery rate (FDR) at a 0.05 level.

| Test | Onset Latency Diff. (P-Value) | Peak Latency Diff. (P-Value) |
| --- | --- | --- |
| (1) > (2) | 0.028* | 0.102 |
| (6) > (7) | 0.034* | 0.645 |
| (8) > (9) | 0.001* | 0.21 |
| (3) > (4) | 0.047* | 0.091 |
| (3) > (5) | 0.003* | 0.091 |
| (4) > (5) | 0.047* | 0.459 |
| (2) ≠ (9) | 0.051 | 0.074 |
| (7) ≠ (9) | 0.051 | 0.006* |
| (2) ≠ (7) | 0.298 | 0.074 |

### 2.3 Emergence of Mirror Invariance

Previous research has shown that the visual system exhibits mirror invariance—the tendency to represent mirrored images similarly across neural populations [5, 14, 16–18]. Here, we tested the hypothesis that mirror invariance constitutes an intermediate computational stage between view-specific and fully view-invariant representations. To examine the temporal dynamics of mirror invariance, we adapted our multivariate pattern analysis approach. In particular, the SVM classifier was trained to pairwise discriminate identity, utilizing train and test sets tailored to detect face representations that were non-invariant, mirror invariant, and fully invariant with respect to head view. For non-invariant representations, classifiers were trained and tested using all four views (left half, left full, right half, right full), with distinct trials randomly selected from each set. For mirror invariance, the classifiers were trained on one of the four views (e.g., left half profile) and tested on its mirrored counterpart (e.g., right half profile). For full invariance, the classifiers were trained on one view and tested on a non-mirrored but different view (e.g., train on left half profile, test on right full profile). To ensure direct comparisons, we excluded the frontal view, and used the same number of trials as in the mirror and fully invariant cases.

The resulting time courses, averaged across participants, identity pairs, and trained views, are shown in Fig. 3. We observed a gradual progression towards increasingly invariant identity representations. Non-invariant representations to head view emerged the earliest at 64 ms (49 - 75 ms), followed by mirror invariant representations at 75 ms (14 - 82 ms), and fully invariant representations emerged last at 89 ms (77 - 98 ms). The onset latency difference between non-invariant and fully invariant representations was statistically significant (Δ = 25 ms, *P* = 0.003*, FDR corrected). We also observed significant differences between non-invariant and mirror-invariant representations (Δ = 11 ms, *P* = 0.047*, FDR corrected), as well as between mirror-invariant and fully invariant representations (Δ = 14 ms, *P* = 0.047*, FDR corrected), although these effects were weaker. Finally, we found no significant differences for peak latencies across conditions. Tables 1 and 2 summarize the onset and peak latencies, along with and corresponding statistical tests for latency differences (conditions 3, 4, and 5).

**Fig. 3:**
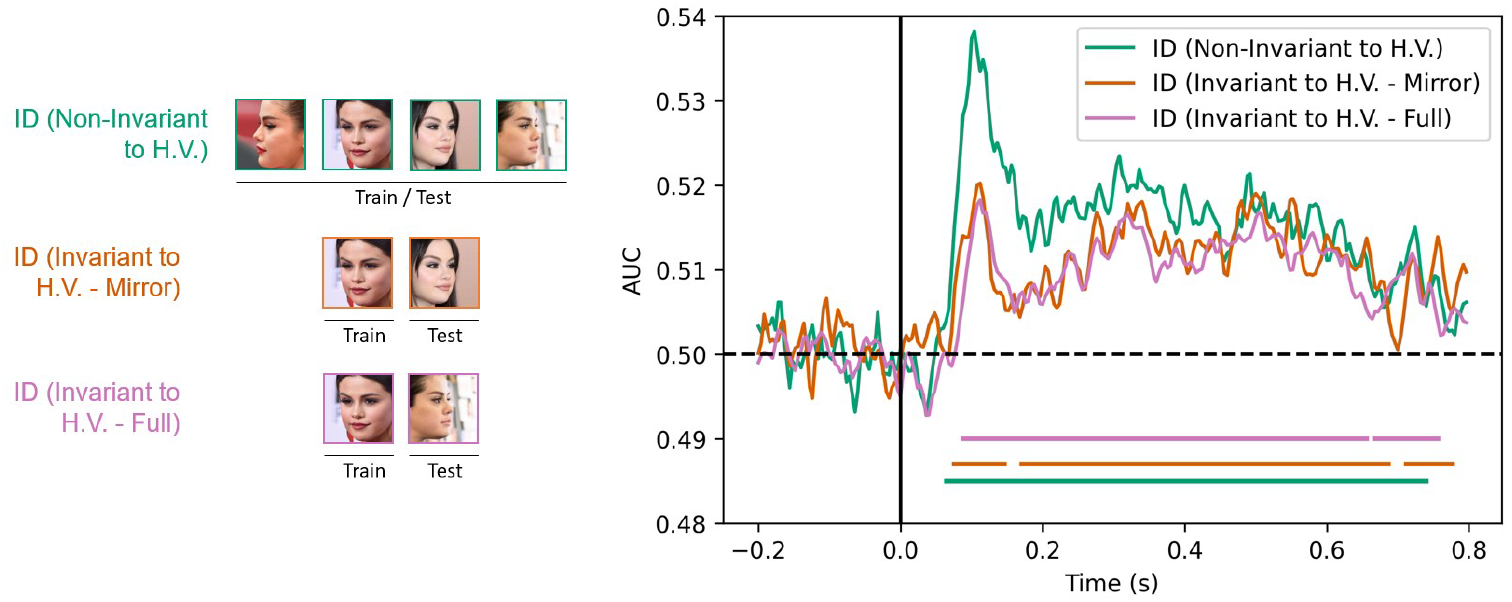
Time course of identity decoding non-invariant, mirror-invariant, and fully invariant to head view. Pictorial representations illustrate cross-decoding approaches for the different cases. Time 0 indicates image onset. Lines below plots indicate significant times using cluster-based sign permutation test (cluster-defining threshold p *<* 0.05, and corrected significance level p *<* 0.05).

These findings support the view that mirror invariance serves as an intermediate stage in the formation of view-invariant face representations [5, 14, 17]. Crucially, our results extend prior work by revealing the full time course from view-specific to fully invariant identity representations, occurring rapidly within a 30 ms window.

### 2.4 Temporal Dynamics of Identity, Age, and Gender Representations

The human brain may extract information about different face dimensions, such as age, gender, and identity, at distinct stages of processing. To investigate this, we performed time-resolved decoding analyses using linear SVM classifiers for each dimension. For age and gender, we conducted binary classifications (young vs. old, female vs. male) using a 6-fold cross-validation procedure, with trials randomly distributed across folds. To assess generalization across identities, we implemented a cross-identity decoding approach by assigning different identities to different folds. This approach allowed us to isolate age- and gender-related information that was invariant to identity. For identity decoding, we used a 5-fold cross-validation scheme with generalization across head views, as illustrated in Fig. 2. Despite the variation in number of folds, due to the constraints of our stimulus set (6 identities, 5 head views), these differences had minimal impact on our main variable of interest: decoding latency.

The time courses of age and gender decoding are depicted in Fig. 4a and b. Visual representations of age emerged at 55 ms (42-80 ms), followed by identity-invariant age representations at 78 ms (54 - 85 ms). This difference in onset was statistically significant (Δ = 23 ms, *P* = 0.034*). Similarly, visual representations of gender first appeared at 75 ms (58-89 ms), followed by identity-invariant gender representations at 91 ms (85-158 ms). The onset latency difference was also statistically significant (Δ = 16 ms, *P* = 0.001*).

**Fig. 4:**
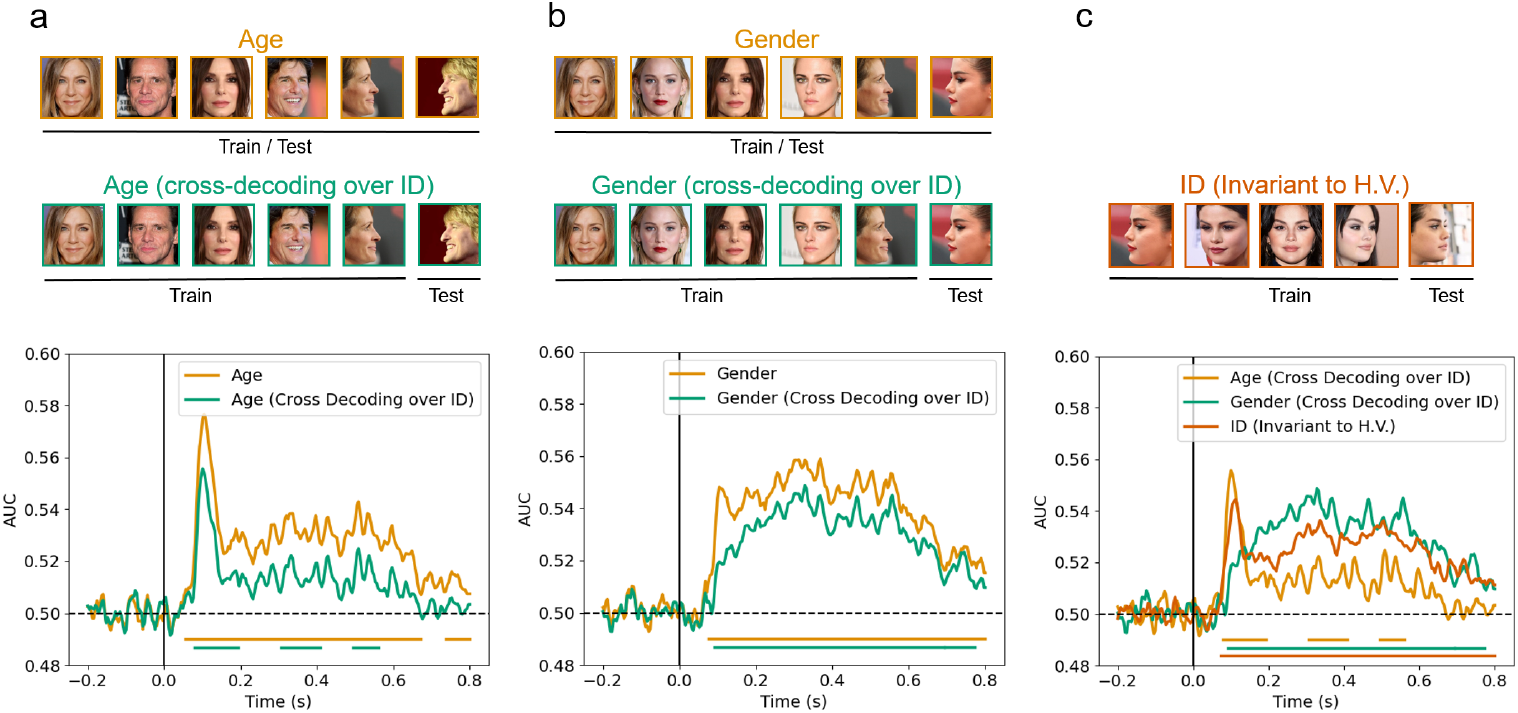
Time courses of age, gender and identity decoding. (a) Temporal dynamics of age decoding. (b) Temporal dynamics of gender decoding. (c) Time courses of age, gender, and identity decoding. Lines below plots indicate significant times using cluster-based sign permutation test (cluster-defining threshold p *<* 0.05, and corrected significance level p *<* 0.05).

The time courses of age, gender, and identity are compared in Fig. 4c. Identity representations invariant to head view emerged first at 73 ms (61-77 ms), followed by age at 78 ms (54-85 ms), and gender 91 ms (85-158 ms). While identity and age appeared to emerge earlier than gender, the difference in onset latency did not reach statistical significance (p = 0.051). Notably, the time course for gender decoding differed qualitatively from those of identity and age. Unlike the latter, which showed early peaks, gender decoding peaked much later at 328 ms (256-556 ms), a latency significantly delayed relative to age (p = 0.006).

In summary, and in contrast to previous findings [11], our results indicate that identity, age, and gender representations emerge at similar times, consistent with partially parallel processing. However, the later peak in gender decoding may reflect more prolonged or distributed processing for this dimension.

### 2.5 Task Effects on Identity, Age, and Gender Representations

To explore the influence of task on face representations, participants completed two distinct tasks in separate sessions: an identity-based one-back task (Task ID) and an image-based one-back task (Task Stim).

Fig. 5 shows the decoding time courses for age, gender, and identity under both task conditions. Identity-invariant age representations emerged at 78 ms (54 - 85 ms) for Task A and 80 ms (59 - 88 ms) for Task B. Identity-invariant gender representations emerged at 91 ms (85 - 158 ms) for Task A and 90 ms (57 - 113 ms) for Task B. Identity representations invariant to head views emerged at 73 ms (61 - 77 ms) for Task A and 78 ms (47 - 82 ms) for Task B. Notably, no statistically significant differences in onset latencies were observed across tasks, suggesting similar early processing dynamics independent of task demands.

**Fig. 5:**
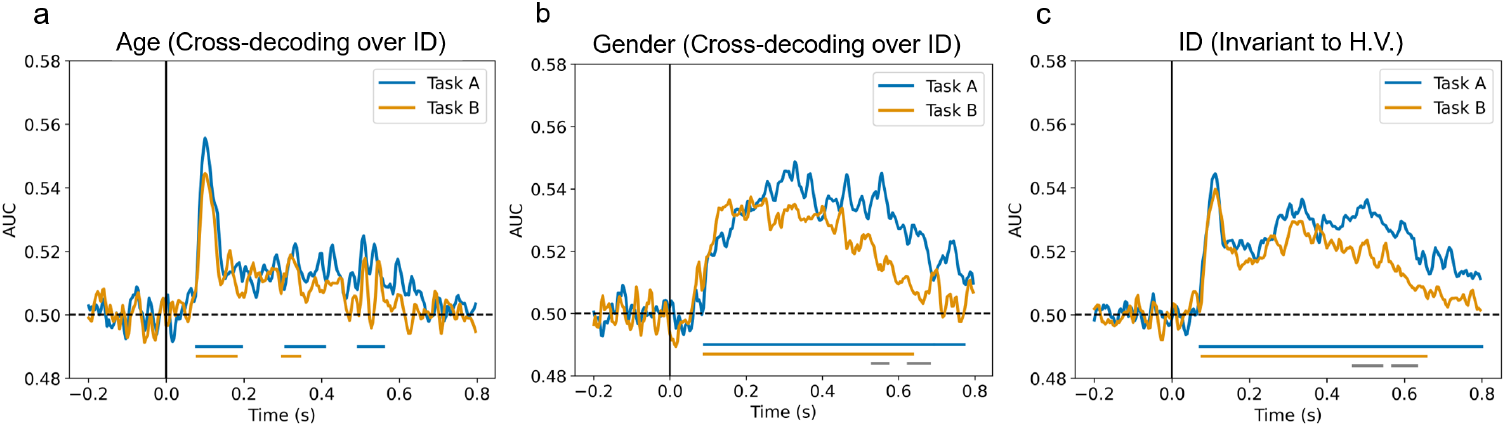
Task effects on identity, age, and gender representations (a) Time courses of age decoding for the two tasks, under cross-decoding over identity. (b) Time courses of gender decoding. (c) Time courses of identity decoding invariant to head view for the two tasks. Lines below plots indicate significant times, with black lines indicating significant differences between the two tasks, determined using cluster-based sign permutation tests (cluster-defining threshold p *<* 0.05, and corrected significance level p *<* 0.05).

Furthermore, we compared the full decoding time courses for AUC differences using cluster-based sign permutation tests. We found no task-related differences for age decoding. However, gender decoding differed between tasks in two late time windows (531 - 572 ms and 625 - 681 ms), and identity decoding differed between 468 - 542 ms and 569 - 632 ms. In both cases, decoding accuracy was higher in Task A than in Task B, suggesting that the more cognitively demanding identity task enhanced later representations of gender and identity.

### 2.6 Source Localization of Identity, Age, and Gender Representations

The human brain possesses a distributed neural system dedicated to face perception, encompassing distinct regions specialized in various facets of face processing. Yet, it remains uncertain whether facial attributes such as identity, age, and gender are processed differently across these regions. To address this question, we localized the activity patterns identified by the SVM classifier during the binary decoding of identity, age, and gender in Task A. We first extracted classifier activation patterns at the early peak decoding latencies: 108 ms for identity, 104 ms for both age and gender. These sensor-level activation patterns—obtained by transforming classifier weights into activation maps—were projected onto individual cortical surfaces and then aligned to a standard brain template (Fig. 6a).

**Fig. 6:**
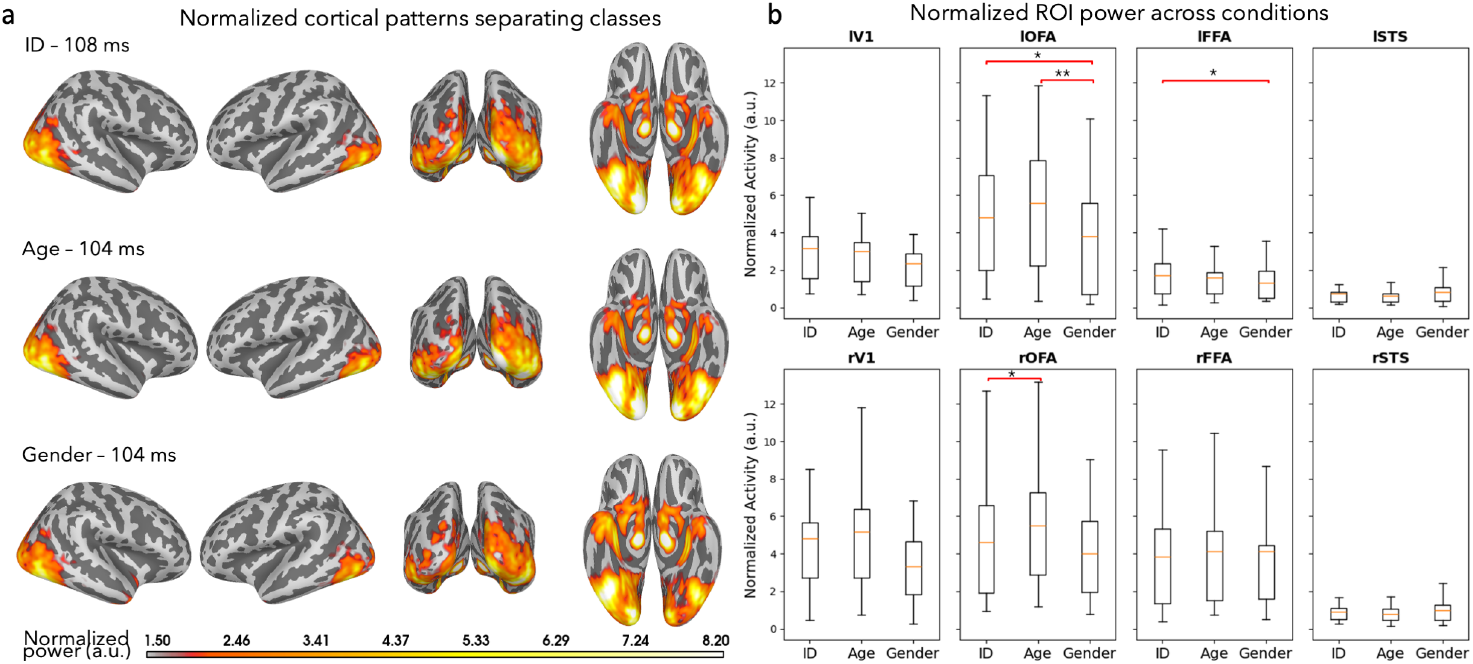
Source localization of identity, age, and gender representations at peak decoding times. (a) Activation maps averaged across subjects, normalized to total unit power across the brain, highlighting the spatial distribution of neural activity. (b) Comparative analysis of activation levels in three face-selective regions of interest (ROIs)—the occipital face area (OFA), fusiform face area (FFA), and posterior superior temporal sulcus (STS). The primary visual cortex (V1) is included as a control region for comparison, providing a comprehensive view of the neural dynamics underlying face attribute processing.

The resulting cortical activation maps, averaged across participants and normalized to total unit power across the entire brain, are displayed in Fig. 6a. Remarkably, these cortical maps showed substantial spatial overlap across the three facial dimensions, suggesting shared early-stage neural substrates for processing identity, age, and gender. To scrutinize more nuanced patterns for subtle differences, we conducted a region-of-interest (ROI) analysis within three canonical face-selective areas: occipital face area (OFA), fusiform face area (FFA), and posterior superior temporal sulcus (STS), as defined by an fMRI-based probabilistic map [19]. Primary visual cortex (V1) serves as a control region (Fig. 6b). Statistical comparison revealed that identity representations were stronger than gender in the left OFA (Δ = 4.96*e −* 05, p = 0.023, FDR corrected), and in the left FFA (Δ = 2.04*e −* 05, p = 0.019, FDR corrected). Age representations also exceeded gender in the left OFA (Δ = 8.60*e −* 05, p = 0.0048, FDR corrected), and identity was stronger than gender in the right OFA (Δ = 4.33*e −* 05, p = 0.019, FDR corrected). These differences are highlighted in red brackets in Fig. 6b.

## 3 Methods

### 3.1 Participants

Nineteen healthy volunteers with normal or corrected-to-normal vision participated in the study. One subject (#14) was excluded due to excessive motion during the recording. Data from 18 subjects (eleven female, seven male) with mean age 25.8 and SD 6.8 remained for the MEG analysis. Of the final sample, 61% identified as White, 17% identified as Asian, 17% identified as Hispanic, and 5% identified as Black. The chosen sample size was based on previous studies using multivariate decoding of EEG/MEG data (e.g. [11]). Sixteen participants were right-handed, while two were left-handed. All subjects provided informed, written consent prior to the experiment. The Massachusetts Institute of Technology (MIT) Committee on the Use of Humans as Experimental Subjects approved the experimental protocol (COUHES No 2105000367) and the study was conducted in compliance with all relevant ethical regulations for work with human participants.

### 3.2 Experimental design and Stimuli

To investigate the temporal dynamics of familiar face processing, subjects viewed face images of famous celebrity identities while monitoring for consecutive repetitions of the same person (i.e., identity based 1-back task; Fig. 1c (left)) and consecutive repetitions of the same image (i.e., image based 1-back task; Fig. 1c (right)) in the MEG. The stimuli used comprised naturalistic images of twelve well-known actors and musicians in the United States. Celebrities were split evenly by gender (male, female) and age (younger, older). Note that here, by gender, we refer to the sex of an individual. For younger individuals, images used in the study were captured when they were under 30 years of age. For older individuals, the images used were taken when they were over 50 years of age.

To ensure that all subjects were in fact familiar with the set of familiar identities, subjects completed an online screening task prior to the study. In this screening, we presented them with one image for each of the 12 identities and asked if they were familiar with the person shown. Only subjects who recognized at least eleven out of twelve familiar identities (e.g., by giving their names or contexts in which they remembered the person) were included in the study. Subjects who missed a single individual were instructed to watch a 10-minute supplementary video about the misidentified celebrity before coming into the lab. Subjects were quizzed on each person upon entering the lab to ensure compliance.

Final stimuli used in the MEG study consisted of fifteen naturalistic color images of each of the 12 identities for a total of 180 stimuli. For each identity, we selected fifteen images from GettyImages or Google search engines which varied in several aspects such as expression (at least five smiling and five neutral facial expressions), five different head orientations (left profile, left half view, direct view, right half view, right profile), eye gaze (for the half views and direct views, at least one image looking directly at the camera and one image with averted gaze were selected), lightning, hair, etc. We limited selection of images with gray hair or facial hair to reduce low level confounds of age and gender. We then standardized all images to a template by rotating, scaling and cropping them based on the position of the nose tip, eyes, and the mouth center. Images shown are not examples of the original stimulus set due to copyright; the exact stimulus set is available at https://osf.io/eh54u.

During the MEG experiment, subjects viewed trials of face images (Fig. 1a). Each trial started with the presentation of a face image for 0.4 s followed by a 0.7–1.55 s interstimulus interval (ISI; uniformly sampled between 0.8 and 1 s) during which a gray screen was presented. Subjects were instructed to respond via button press to a consecutive repetition of an identity and identical image during image presentation or during ISI based on the task. To avoid artifacts due to eye movements or blinking, subjects were instructed to fixate a black fixation cross in the center of the screen during image presentation (i.e., presented between the tip of the nose and the eyes of a face) and ISI. They were further asked to blink at the same time when giving a button response, as these trials were not included in the data analysis.

Subjects viewed 20 blocks of trials in which each of the 90 images were presented once randomly interleaved with 22 task trials (1-back task) per block for a total of 440 trials. Stimulus presentation was controlled and responses collected using Psychtoolbox 3 for Matlab [20]. The experiment lasted around 60 min.

### 3.3 MEG Recording and Pre-processing

Each of the 18 participants completed two MEG sessions, each consisting of image and identity-based 1-back tasks. The task order was counterbalanced, with approximately half of the participants starting with Task A, and the remaining participants initiating with Task B. During each session, participants completed three stages: (i) an eye-tracker calibration, (ii) a static localizer, and (iii) the main task involving face images. In the eye-tracker calibration (i), participants were instructed to follow a dot with their eyes on the screen to calibrate the eye-tracking device. Following that, in the static localizer stage (ii), grayscale images of faces and objects were presented, and participants were instructed to respond via button press to consecutive repetitions of an image. A total of 360 trials, including 60 task trials and 300 trials (10 repetitions of unique 15 faces and 15 object images), were collected during the static localizer. Each trial consisting of the presentation of an image for 0.4 seconds, followed by a 0.2-0.3s interstimulus interval featuring a gray screen. A 0.75s buffer was included if the presented image corresponded to a task trial, and 0.5s if it was not a task trial. An additional 0.5s buffer was added if participants pressed the button. Consequently, the total interstimulus interval could range from 0.7 to 1.55s. The last stage in the MEG experiment was the main task.

MEG data were collected using a 306-channel Elekta Triux system with a 1000 Hz sampling rate, and were filtered online between 0.01 and 330 Hz. The position of the 12 head was tracked during MEG recording based on a set of five head position indicator coils placed on particular landmarks on the head. We preprocessed the raw data with Maxfilter software (Elekta, Stockholm) to remove head motion and to denoise the data. The “-movecomp” parameter within Maxfilter facilitated the correction of head movements that may occur during the MEG recording session. This correction ensures the alignment of MEG data across different time points. Additionally, “-tSSS” temporal signal space separation technique employed to mitigate environmental noise and magnetic interference, was applied. tSSS separates signals originating from the brain from those stemming from external sources, contributing to a cleaner and more reliable dataset.

We utilized Python (Spyder IDE version 5.4.2) to extract epochs ranging from −200 to 800 ms relative to image onset. For our magnetometers, we employed a flat criterion of 1 fT and for gradiometers, a flat criterion of 1 fT/cm, which establishes the minimum acceptable peak-to-peak amplitudes for each channel type in an epoch. Furthermore, we implemented a rejection threshold of 8000 fT for magnetometers and 8000 fT/cm for gradiometers, setting the maximum acceptable peak-to-peak amplitudes for each channel type in an epoch. To mitigate Electrooculogram (EOG) artifacts, we applied EOG projectors. Subsequently, each epoch underwent baseline correction by removing the mean activation from each MEG sensor between −200 ms and stimulus onset. The epochs were then low-pass filtered to 35 Hz. Additionally, we computed and saved the noise covariance.

The initial MEG data, recorded at a high sampling rate of 1000Hz, underwent a purposeful downsampling to 250Hz. This downsampling step was implemented to enhance computational efficiency during subsequent machine learning decoding analysis. Following the decoding analysis, the dataset was upsampled back to its original 1000Hz sampling rate to compute significant clusters, peak, onset time and confidence interval. Upsampling back to 1000Hz provides higher precision in the temporal domain, allowing for a more accurate characterization of the temporal properties of the MEG signal.

### 3.4 MEG temporal decoding analysis

We used multivariate pattern analysis to extract temporal information about the face stimuli from the MEG data. To obtain the time course with which experimental conditions, i.e. age, gender and identity, are discriminated by MEG, we used linear support vector machine (SVM) classification [21], as implemented in the libsvm software [22] with a fixed regularization parameter C = 1.

For each participant, we performed temporal decoding of experimental conditions without generalization by initially selecting epochs from the complete MEG epoched data that aligned with specific pairwise conditions. For instance, in the age experimental condition, the pairwise combination comprised epochs categorized as young and old. This process was repeated for all pairwise combinations, employing a five-fold cross-validation where we tested on one group and trained on the remaining groups. The outcome was an average decoding accuracy value per subject and time point.

When extending temporal decoding to involve generalization from one experimental condition (Condition 1) to another (Condition 2), a specialized training and testing set were designed. The model was trained on a subset of Condition 2 corresponding to Condition 1 and tested on the remaining part of Condition 2. This approach allowed us to generalize temporal decoding patterns from one experimental condition to another, providing a comprehensive understanding of information processing across different conditions.

### 3.5 Source reconstruction of neural activity underlying classification

We computed the AUC over time from sensor-level classification results for Identity, Age, and Gender previously described in section 3.4 (non-invariant cases only). Subsequently, we determined the time of peak AUC for each condition, which will be used later in the following source-level analysis. To investigate the cortical patterns of activity underlying the classification, we first transformed sensor-level classifier weights using a method described by Haufe [23]. Next, source reconstruction was performed using a minimum-norm estimate (MNE) (Gramfort, 2013; Hämäläinen & Ilmoniemi, 1994) distributed model containing 20484 dipolar sources, producing cortical activity maps specific to individual anatomy which were then morphed to the Freesurfer standard brain (fsaverage). Raw classifier weights are difficult to interpret as high values may result from sensor noise removal rather than meaningful task-related signals [23, 24]. Therefore, transforming these classifier weights into neural activity using the method by Haufe would enhance their interpretability. The general steps for performing this transformation followed by source reconstruction are also outlined in the MNE-python tutorial online under the section “Decoding MVPA”. For comparability across subjects, regions, and conditions, the activity at each vertex was squared, yielding vertex power, and spatially normalized by dividing it by the average unit power per vertex, resulting in a cortical map showing the proportion of total power activated per vertex for each condition (Fig. 6a). Using a functional MRI-based probability map previously developed by the Kanwisher lab [19], we localized three face-selective regions-of-interest (ROIs) – the occipital face area (OFA), fusiform face area (FFA), and posterior superior temporal sulcus (STS) – in each hemisphere of the Freesurfer standard brain. Additionally, we defined the primary visual cortex (v1) as a reference using the Desikan-Killiany anatomical atlas [25] provided by Freesurfer (see Fig. 7). For each of the ROIs, we computed the mean normalized power across vertices within the ROI at the time of peak AUC obtained earlier. To assess differences in mean normalized power between conditions at each ROI, we performed a two-tailed permutation test (9999 permutations) at a critical level of 0.05 with FDR-correction across 24 comparisons using the Benjamini and Hochberg procedure [26].

**Fig. 7:**
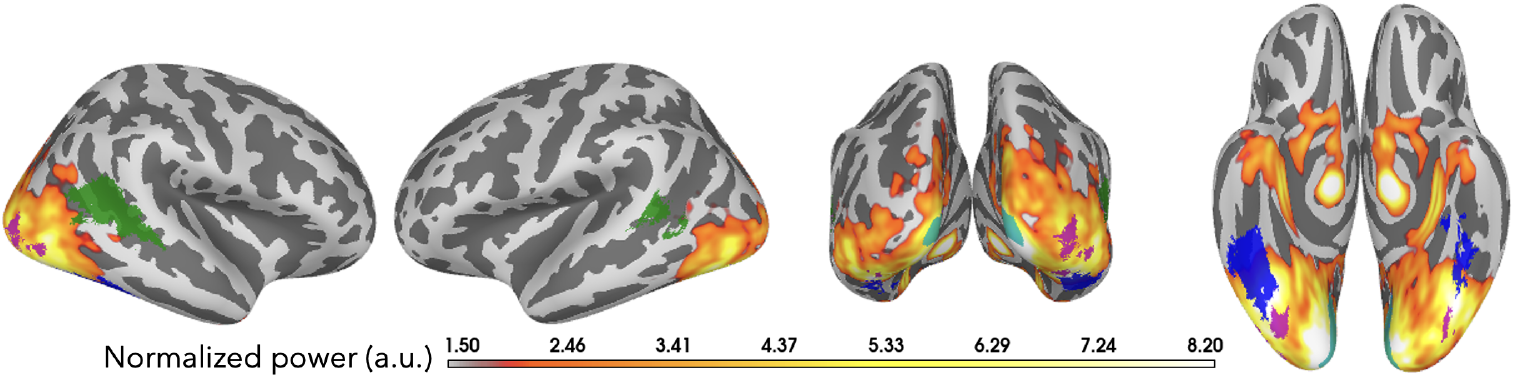
Regions-of-interest displayed over the standard brain and typical cortical activity. Colored patches highlight the occipital face area (magenta) fusiform face area (blue), posterior superior temporal sulcus (green), and primary visual area (cyan). Orientation views (from left to right): right lateral, left lateral, posterior, anterior.

### 3.6 Onset and peak latency analysis

To test for statistical differences in onset or peak latencies between different face dimensions, we performed bootstrap tests. We bootstrapped the subject-specific decoding accuracy 1000 times to obtain an empirical distribution of the mean onset (i.e., earliest significant time point post stimulus onset) and peak latencies (i.e., significant time point at maximum peak post stimulus onset) across each bootstrapped group of subjects. We restricted the time window for the peak analysis to 180 ms post stimulus onset, since we were interested in the first peak occurring after stimulus onset, unconfounded from later peaks (e.g., due to stimulus offset responses). The 2.5th and the 97.5th percentile of these distributions defined the 95% confidence interval for onset and peak latency, respectively. For differences between latencies, we computed 1000 bootstrap samples of the difference between two latencies (e.g., onset and peak) resulting in an empirical distribution of latency differences. The number of differences that were equal and smaller (or larger) than zero divided by the number of permutations defined the p-value (i.e., one-sided testing). For a two-sided test these p-values were multiplied by 2. These pvalues were corrected for multiple comparisons using false discovery rate (FDR) at a 0.05 level.

## 4 Discussion

This study examined the spatiotemporal dynamics of familiar face processing in the human brain using time-resolved decoding of MEG signals. We addressed three core questions: (1) Does facial identity become increasingly view-invariant over the course of neural processing? (2) How do the temporal dynamics of identity compare to those of age and gender representations? (3) How do task demands modulate these processes? We found that identity representations emerged rapidly and became progressively tolerant to changes in head view, with decoding first evident for identical views at 64 ms, followed by mirror-symmetric views at 75 ms, and full view-invariant representations at 89 ms. Age and gender representations also emerged early and became increasingly invariant to identity. Finally, later representations of identity and gender were enhanced under an identity-focused task relative to a low-level image-matching task. Source localization analyses confirmed activation in known face-selective regions, including the OFA and FFA, and revealed age-related signals in these areas as well.

Our finding of progressive view invariance for identity aligns with prior reports in both object and face processing. For instance, Kietzmann et al. (2017) [6] and Guntupalli et al. (2017) [3] observed view-tolerant face representations in EEG and fMRI using unfamiliar faces. Our results extend these findings to highly familiar identities and demonstrate that mirror-invariant representations precede full view invariance in time, supporting the proposed hierarchical progression of visual tolerance [14, 18]. Notably, our observed latency for peak view-invariant identity decoding (112 ms) is substantially earlier than those reported in EEG studies using unfamiliar or standardized faces, such as the 250 ms peak observed by Caharel et al. (2015) [7]. This timing difference may reflect the prioritized and more efficient processing of familiar faces [10, 27], or it may result from the use of naturalistic stimuli that retain low-level cues (e.g., hair, clothing) known to aid recognition [9].

We also investigated the relative timing of identity, age, and gender decoding. Identity decoding invariant to head view emerged earliest at 73 ms, followed closely by age at 78 ms and then gender at 91 ms, though differences in onset were only marginally significant. This pattern contrasts with some previous reports. For example, Dobs et al. (2019) [11] and Kaiser et al. (2020) [28] suggested a coarse-to-fine sequence in which basic attributes like gender and age are extracted before identity. One plausible explanation for our different ordering is the use of highly familiar faces in our study, which may have led to faster engagement of person-specific memory systems. Indeed, Ambrus et al. (2019) [27] also observed early identity decoding for familiar faces using naturalistic stimuli. Another factor could be our use of color images with rich contextual cues, which may facilitate identity recognition more than grayscale or controlled face stimuli used in other studies. Thus, the temporal hierarchy of face attribute decoding may depend on both familiarity and the ecological validity of the stimuli.

Importantly, our results suggest that age and gender information, like identity, becomes increasingly invariant across other dimensions. We found that both age and gender representations generalized across identity with only modest temporal delay. This finding expands prior work showing that facial age and gender can be decoded from MEG signals [15, 29], and it supports models in which invariant representations of these attributes are formed through dynamic neural processes [30]. To our knowledge, this is the first study to demonstrate increasing identity-invariance of age and gender representations over time.

Task demands also modulated facial processing. While early latencies were not significantly affected by task type, later identity and gender representations were stronger during the identity-based task. This is consistent with theories of top-down modulation in face processing [10], and with prior findings that task relevance enhances representational strength [11]. Our results support the view that early processing stages are largely automatic, while later stages reflect strategic engagement based on task demands.

We also leveraged MEG source localization to examine the spatial sources of decoded representations. Identity, age, and gender signals all showed consistent activation in the right-lateralized face processing network, especially the OFA and FFA. Our results align with prior fMRI studies identifying the FFA and anterior temporal cortex as core regions for identity representation [4, 31, 32], and studies that localize gender information to OFA and FFA but not STS [29]. Notably, we provide one of the first demonstrations of robust age-related decoding localized to FFA and OFA using MEG source estimation.

Despite these strengths, our study has several limitations. First, although our use of naturalistic, familiar images enhances ecological validity, it introduces uncontrolled variability in low-level features, such as background and lighting, which could partially drive decoding. Second, although decoding performance was statistically robust, effect sizes were modest, and classifier performance remained relatively close to chance, suggesting these effects should be interpreted cautiously. Third, familiarity in our study was based on media exposure, not personal relationships. Prior work has shown stronger neural representations for personally familiar faces [27], which may limit generalizability to real-life interactions. Lastly, while source localization was informed by classifier weights, the interpretation of these maps depends on assumptions that can be violated in complex sensor-level transformations [23].

In conclusion, our study provides new evidence that identity representations of familiar faces become increasingly view-invariant over time and that age and gender attributes also exhibit generalization across identity. These findings refine our understanding of the temporal and spatial architecture of face perception and emphasize the importance of task and familiarity in shaping these processes. Future work should explore how these dynamics vary with personal vs. media familiarity and investigate their neural substrates using combined MEG-fMRI or intracranial methods. These insights may also inform the design of computational models that aim to emulate human-like robustness in face recognition across changing views and identities.

## 5 Conclusion

This study provides new insights into the spatiotemporal dynamics of familiar face processing, highlighting how the brain extracts identity, age, and gender information from naturalistic face images across varying head orientations. Using MEG and time-resolved decoding, we uncovered distinct temporal profiles for each attribute and demonstrated how these representations evolve toward greater invariance during visual processing.

Critically, we found that facial identity representations become progressively view-invariant over time, with mirror symmetry emerging before full view invariance—supporting a hierarchical model of visual generalization. Age and gender information also emerged early and became increasingly invariant to identity, suggesting shared computational principles across facial dimensions. While early latencies were largely unaffected by task, later stages of identity and gender processing were enhanced under identity-focused attention, revealing a dynamic interplay between bottom-up and top-down mechanisms. Behavioral data further confirmed that identity-based tasks imposed higher attentional demands.

Together, these findings advance our understanding of how the brain recognizes familiar faces under real-world variability. By combining high-temporal-resolution MEG with multivariate decoding and source localization, this work provides a temporally precise account of how invariant face representations emerge in the human brain. These results inform computational models of face recognition and offer valuable guidance for designing neurotechnologies in fields such as biometrics, social robotics, and clinical assessment of face perception deficits.

## Acknowledgments

This work was supported by the National Eye Institute of the NIH under award number 1R01EY033638 (to DP), the United States-Israel Binational Science Foundation grant 2020805 (to AA), and the National Institute on Aging of the NIH under award numbers RF1AG074204 and RF1AG079324 (to DP). The content is solely the responsibility of the authors and does not necessarily represent the official views of the National Institutes of Health. The authors have no relevant personal financial or non-financial interests to disclose.

